# Quantification of nuclear protein dynamics reveals chromatin remodeling during acute protein degradation

**DOI:** 10.1101/345686

**Authors:** Alexander J. Federation, Vivek Nandakumar, Hao Wang, Brian C. Searle, Lindsay K. Pino, Gennifer Merrihew, Ying S. Ting, Nicholas Howard, Tanya Kutyavin, Michael J. MacCoss, John A. Stamatoyannopoulos

## Abstract

Sequencing-based technologies cannot measure post-transcriptional dynamics of the nuclear proteome, but unbiased mass-spectrometry measurements of chromatin-associated proteins remain difficult. In this work, we have combined facile nuclear sub-fractionation approaches with data-independent acquisition mass spectrometry to improve detection and quantification of nuclear proteins in human cells and tissues. Nuclei are isolated and subjected to a series of extraction conditions that enrich for nucleoplasm, euchromatin, heterochromatin and nuclear-membrane associated proteins. Using this approach, we can measure peptides from over 70% of the expressed nuclear proteome. As we are physically separating chromatin compartments prior to analysis, proteins can be assigned into functional chromatin environments to illuminate systems-wide nuclear protein dynamics. The integrity of nuclear sub-compartments were validated with immunofluorescence, which confirms the presence of key markers during chromatin extraction. We then apply this method to study the nuclear proteome-wide response to pharmacological degradation of the BET bromodomain proteins. BET degradation leads to widespread changes in chromatin composition, and we discover global HDAC1/2-mediated remodeling of chromatin previously bound by BET bromodomains. In summary, we have developed a technology for reproducible, comprehensive characterization of the nuclear proteome to observe the systems-wide nuclear protein dynamics.

## Main Text

Regulatory DNA is characterized by the cooperative binding of sequence-specific transcription factors (TFs) and their associated cofactors that regulate local chromatin structure and nearby gene expression [1-4]. While genome-wide catalogues of DNA harboring regulatory potential are nearing saturation [5,6], comprehensive characterization of the protein compartment responsible for transcriptional regulation remains in its infancy [7]. There are more than 1,500 transcription factors, cofactors, and chromatin regulators encoded in the mammalian genome, and we have limited understanding of the genomic occupancy and contribution to chromatin regulation for most of these [8]. In many studies, mRNA levels or promoter chromatin state are used as a proxy for the activity of a gene product [9,10], but these measurements do not correlate highly with protein abundance or activity. Furthermore, these methods do not capture subcellular location or proteoform diversity. Chromatin immunporecipitation (ChIP) and other DNA-centric assays are commonly used to measure nuclear protein dynamics, however, these approaches are also inherently limited. DNA-centric approaches require specific antibodies or protein tagging for the protein of interest and, most importantly, they can only assay a single protein per experiment, making systems-level measurements of chromatin dynamics impossible.

Mass spectrometry-based proteomics can be used to measure systems-level protein dynamics in an unbiased way, but fully characterizing the nuclear proteome by mass spectrometry is nontrivial. Many chromatin proteins are expressed transiently, at low levels, or are difficult to extract from the nucleus [11-13]. Previous attempts to fractionate or enrich for the nuclear proteome exhibit detection rates ranging from 10%-40% of expected TFs and cofactors [14-20]. Moreover, sample preparation for these methods often include laborious protein fractionation strategies, limiting scalability and widespread adaptability [19, 20].

Besides low TF abundance, additional difficulties arise from high concentrations of structural proteins in the nucleus, which suppress detection of transcriptional regulators in typical nuclear extract proteomics. Data-dependent acquisition (DDA) mass spectrometry approaches fragment and sequence peptides from the most abundant chromatography peaks, leading to infrequent and stochastic detection of lowly abundant peptides. This leads to incomplete sampling in large-scale datasets containing many samples, making it impossible to distinguish between missing measurements due to stochastic detection verses the true absence of a peptide of interest [21].

To comprehensively quantify protein dynamics across various chromatin environments, we developed Chromatin Enrichment by Salt Separation followed by Data-Independent Acquisition (ChESS-DIA) (Supplementary Figure 1). Protein-nucleic acid interactions are largely driven by electrostatic attraction [22], and by treating isolated nuclei with increasing salt concentrations, we observe a bimodal disruption and release of proteins from surrounding nuclear material (Supplementary Figure 2). As previously reported, this bimodality reflects differences in the physical characteristics of open regions of “active” euchromatin versus “closed” heterochromatin in the nucleus [22]. We leverage these differences to combine a facile, scalable nuclear separation protocol followed by label-free data-independent mass spectrometry acquisition, allowing the comprehensive isolation and chromatin environment assignment for greater than 70% of the proteins predicted to be expressed in the nuclear proteome. In short, nuclei are isolated from cells or tissue of interest, then subjected to a series of extraction conditions that isolate (1) unbound proteins in the nucleoplasm, (2) proteins bound in accessible euchromatin, (3) bound in heterochromatin and (4) insoluble structural proteins. This insoluble fraction, which primarily contains histones and lamina-associated proteins, accounts for a large fraction of the ions measured in a typical nuclear extraction, so their separation from functional chromatin compartments allows for more comprehensive sampling of proteins in euchromatin and heterochromatin. After protein extraction, digestion and desalting, mass spectrometry data is acquired and quantified using on-column chromatogram libraries to increase sensitivity and reproducibility across large datasets [23] (Supplementary Figure 1).

We compared ChESS-DIA with a recently published high-quality nuclear proteomics dataset in an acute myeloid leukemia cell line that utilizes TMT-isobaric mass tagging [24]. Overall, ChESS-DIA detected 33% more TFs and cofactors, and this improvement is dependent on TF family. For example, important developmental TF families that tend to be lowly expressed, such as homeodomains and forkhead proteins, are detected at a 2-fold and 1.8-fold higher rate, respectively. Proteins detected by ChESS-DIA had good peptide coverage, with a median value of 5.5 peptides per protein (Figure 1C). Peptide detection rates generally correlated with protein expression level, and <2% of peptides were detected for genes unlikely to be expressed (Figure 1D, RNA-seq FPKM < 1.0). Gene Ontology (GO) enrichment analysis of the observed proteins show high enrichment scores for annotations related to nucleic acid binding, chromatin regulation and nuclear localization. Among the chromatin compartments, euchromatin shows the strongest GO enrichment (Supplementary Figure 3).

**Figure 1:**
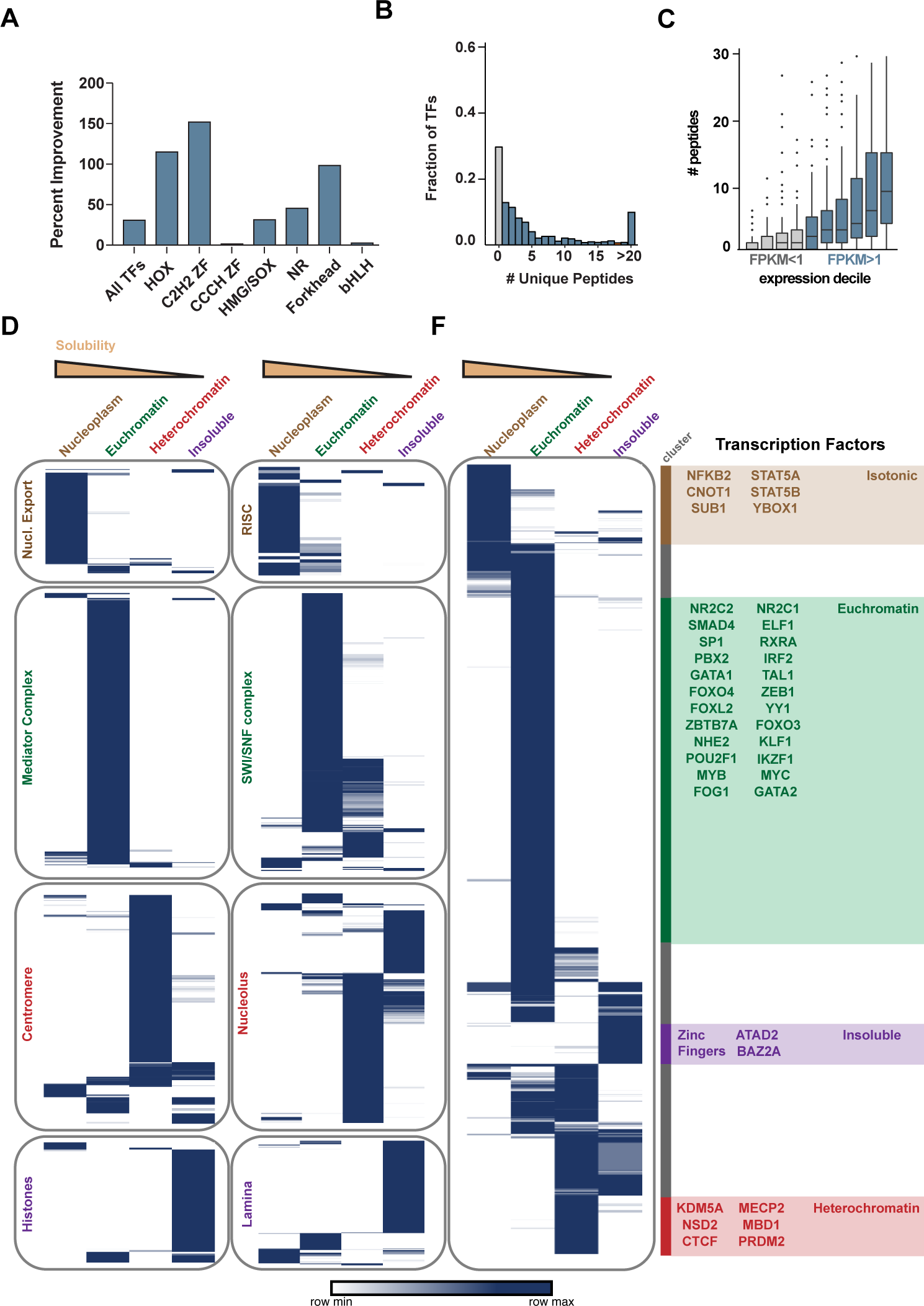
ChESS DIA detection rates for nuclear proteins and assignment into functional chromatin compartments. (A) Protein detection rates in ChESS-DIA compared to Winter et al (2017). (B) Peptides detected per protein for all expressed transcriptional regulatory proteins (C) Peptides detected per protein for different levels of RNA expression by RNA-seq. (D) Peptide quantification for known protein families and complexes. Rows repreent individual peptides and each row is normalized to the max value. (E) Peptide quantification for all TF-associated peptides.

To further validate chromatin compartments extracted with this method, we observed the distributions of several protein complexes with well-established roles in a defined chromatin context (Figure 1D). The exportin family of proteins that regulate nuclear export and the RISC RNA-processing complex are highly enriched in the nucleoplasm, consistent with their nonchromatin related function. The transcriptional coactivating mediator complex is found predominantly in the euchromatin fraction [25], as is the SWI/SNF chromatin remodeler. The SWI/SNF complex can facilitate the transition of polycomb-repressed regions into accessible euchromatin, and interestingly, subsets of the SWI/SNF associated peptides are also found to occupy heterochromatin compartments by ChESS-DIA, particularly peptides mapping to the catalytic subunits SMARCA2 and SMARCA4 [26]. Centromeres, a canonical heterochromatin structure, are highly enriched in this fraction, as are proteins associated with the nucleolus. Histones and nuclear lamina are predominantly insoluble, with the exception of LAP2A, which is a lamina-family protein with an established role regulating gene expression in euchromatin [27].

TFs are found to occupy all four fractions, as expected by their wide range of functionalities within the nucleus (Figure 1E). Nucleoplasmic TFs include NFkB, which is sequestered from chromatin until an activating phosphorylation event. The euchromatic TFs include many lineage-specific and oncogenic TFs that known to dominate the transcriptional regulatory program in K562 cells [28]. Heterochromatic TF include proteins with known roles in nuclear organization (CTCF), interaction with methylated DNA (MECP2, MBD1) and chromatin compaction [29]. Insoluble TFs notably include many zinc finger TFs that constitutively suppress retrotransposon activity [30].

We validated the ChESS-DIA quantification using cellular immunofluorescence (IF) imaging as an orthogonal method of protein quantification. Analysis of extraction kinetics for the euchromatic GATA1 TF shows rapid and complete removal from nuclei after a 5-minute extraction (Figure 2A), quantified across 100 cells per time point (Figure 2B). All TFs studied were completely extracted after 10 minutes (data not shown). All subsequent proteomics data presented in this study employ 20 minute extractions to account for the potential variability in proteome-wide extraction kinetics. IF was then performed on canonical members of each chromatin compartment assayed (Figure 2C). Single-cell IF quantification of TF extractions show a striking correlation with ChESS-DIA based quantification across all fractions. This is even reflected for the BRD4 protein is present in both the euchromatin and heterochromatin fraction, which is reflected in the IF-based quantification. Furthermore, we integrated ChESS-DIA compartment assignments with the large-scale effort to map subcellular locations by IF performed by the Human Protein Atlas (HPA) [31]. The three chromatin-bound fractions, as defined by ChESS-DIA, are only enriched in nuclear-related subcellular locations by IF. ChESS-DIA euchromatin proteins are most likely to be assigned within nuclear speckles, which contain splicing factors and are involved in active transcription within the nucleus [31]. ChESS-DIA insoluble proteins, on the other hand, are most likely to be found associated with the nuclear membrane, consistent with the role of membrane sequestration in gene silencing [31].

**Figure 2:**
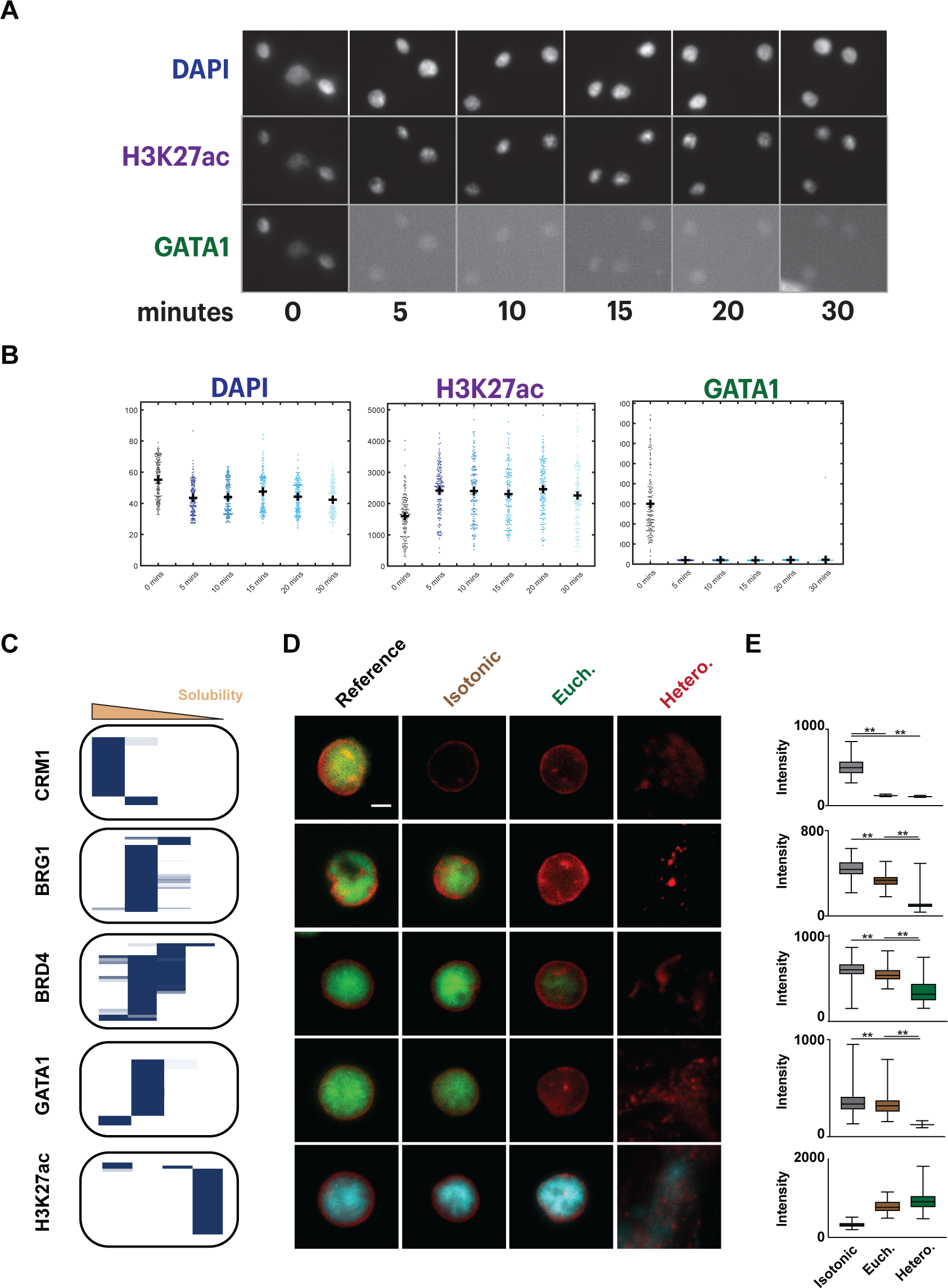
Immunofluoresence reveals extraction kinetics and validates ChESS-DIA measurements. (A) K562 nuclei were treated with euchromatin extraction buffer, then fixed after the indicated time, ranging from 0-30 mins. Fixed nuclei were stained with DAPI and antibodies recognizing H3K27ac and GATA1 and imaged. (B) Immunofluoresence-based quantification for 100 cells per time point. (C) ChESS-DIA distibutions of CRM1, BRG1, BRD4, GATA1, H3K27ac. (D) epresentative single cell images of K562 cell nuclei treated with various extraction buffers for 5’ and immunola-beled for the protein of interest in ‘C. Protein staining in green (CRM1/BRG1/BRD4/GA-TA1) or cyan (H3K27ac). Lamin B1 staining is in red. Scale bar = 5 microns. (E). Legend for ‘E’: Nuclear protein abundance for the various extraction conditions. p < 0.01**

With a validated method in hand, we turned to address a longstanding problem in the pharmacodynamics of transcriptional regulators. Transcriptional regulatory proteins that lack a catalytic domain are often difficult to drug with small molecule inhibitors, but the development of heterobifunctional small molecule compounds that can bind and degrade their target protein effectively target this protein class [24,33]. The primary pharmacodynamic effect of these compounds is a global reorganization of chromatin, but this is difficult to measure with ChIP-based methods or existing nuclear proteomics methods, obfuscating the full effects of these compounds. Therefore, we applied ChESS-DIA to characterize the global chromatin response to acute degradation of the first reported pharmacological degraders that target the BET bromodomains.

The BET bromodomain family contains BRD2, BRD3, BRD4 and BRDT, all which contain tandem bromodomains that function to bind acetylated lysine residues on histones and transcription factors [34,35]. They are expressed constitutively across all tissues and they play important roles in gene regulation and nuclear organization [36]. Furthermore, their function is implicated in many models of human cancer and drugs targeting BETs are currently in clinical investigation for various tumor types [37]. In steady-state cells, BRD2, BRD3 and BRD4 contain partially overlapping but distinct chromatin compartment distributions (Figure 3A), consistent with their partially overlapping but distinct roles in transcriptional regulation [38-40].

**Figure 3:**
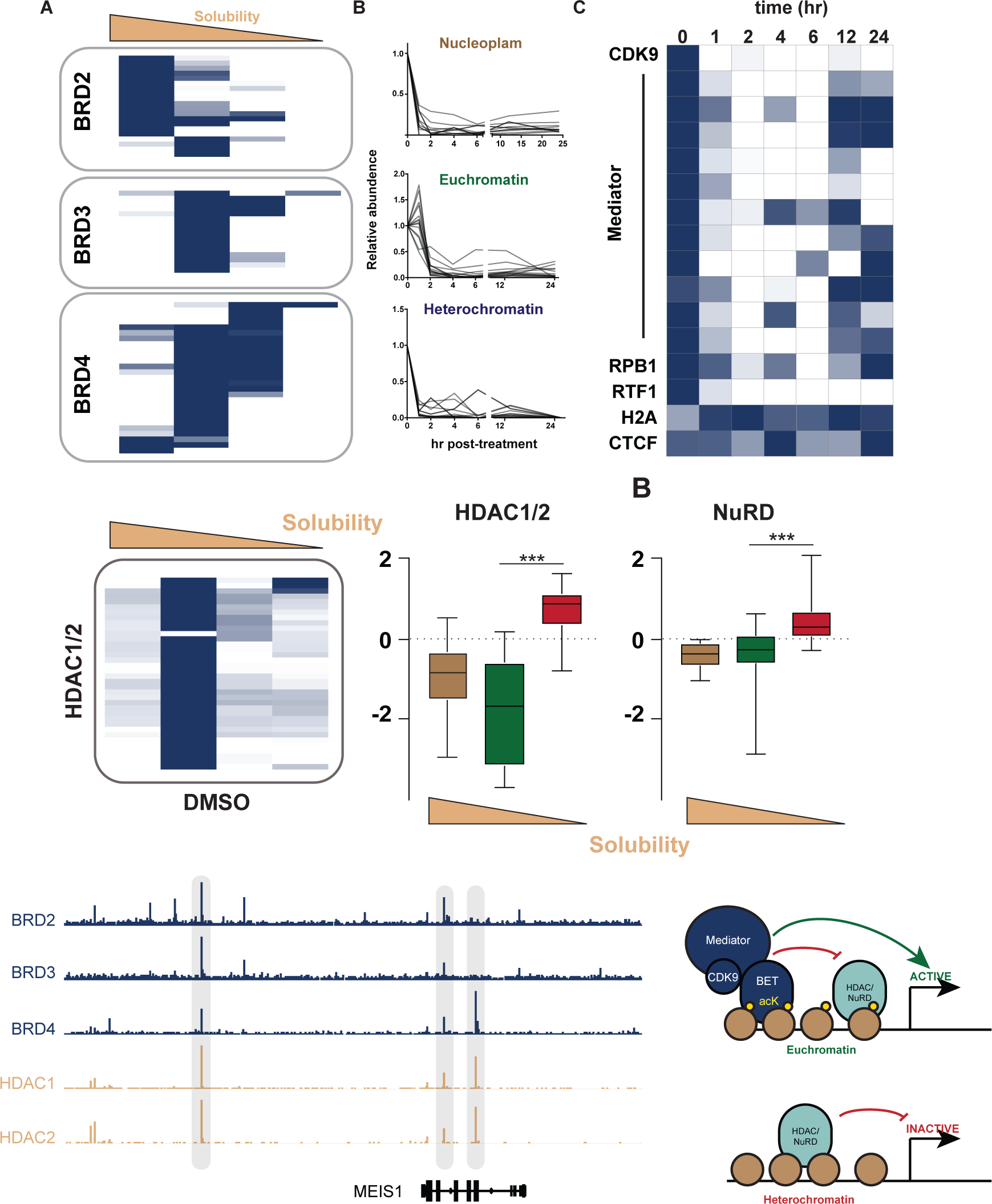
ChESS-DIA reveals chromatin reorginization during acute BET degreda-tion. (A) ChESS-DIA distrubutions of the BET bromodomains in MOLM-13 (B) Response of quantificaions for all BET proteins in each chromatin fraction to **dBET1** time course (C) Response to **dBET1** of peptide quantificaions for BET-interaction partners and control proteins (H2A, CTCF) (D) ChESS-DIA distrubutions of the HDAC1 and HDAC2 in MOLM-13. Fold change of HDAC1/2 and NuRD complex members in response to **dBET1** at 4h (E) Browser track of Cut&Run data normalized by reads per million (F) Schematic showing effects of HDAC activity after **dBET1** treatment

Recently, **dBET1** was developed as a first-in-class small molecule BET bromodomain degradation compound [24]. **dBET1** functions by selectively binding the BET bromodomain and recruits the cereblon-CUL4 ubiquitin ligase complex, leading to BET ubiquitination and proteasome-mediated degradation. In agreement with previously published measurements of **dBET1** kinetics [32], ChESS-DIA confirmed the depletion of total BRD2, BRD3 and BRD4 from all chromatin compartments within 2 hours of treatment with **dBET1** in MOLM-13 acute myeloid leukemia cells. BRDT is a testes-specific protein and is not expressed in these cells. We then looked specifically at the euchromatin protein fraction, where canonical BET bromodomain cofactors are observed. During **dBET1** treatment, CDK9 (pTEFb) and the mediator complex are lost from euchromatin, consistent with previous studies showing loss of these transcriptional cofactors upon loss of BET bromodomains from chromatin as measured by ChIP-seq [41]. Protein subunits of RNA PolII are also depleted from active chromatin, consistent with a global loss in transcriptional activity (Figure 3C) [42].

Looking across all fractions, we observed a striking reorganization of HDAC1 and HDAC2 peptides from euchromatin into heterochromatin with the same kinetics as BET bromodomain loss. HDAC1/2 cooperate as part of the NuRD complex to regulate transcription by deacetylating acetylated histones and promoting chromatin compaction, particularly at bivalent genes during development [43]. Peptides for the other NuRD complex members show the same pattern of euchromatin to heterochromatin transition (Figure 3E), so we hypothesized that this phenomenon was due to chromatin compaction at sites shared by HDAC1/2 and BETs. Indeed, like the BET bromodomains and their cofactors, HDAC1/2 are found predominantly euchromatin (Figure 3F). To test for genome-wide co-occupancy of HDAC1/2 and BET bromodomains, we used Cut&Run to map binding locations of all factors [44]. Indeed, HDAC1/2 Cut&Run peaks display significant overlap with BRD2 (18% of HDAC1/2 shared peaks), BRD3 (5%) and BRD4 (11%). These findings support a model where at steady-state, HDAC1/2 activity is suppressed at active DNA regulatory elements by BET bromodomains. When the BETs are degraded by **dBET1**, HDAC1/2 complexes remaining at regulatory DNA can deacetylate nearby histones, leading to compaction local chromatin after BET loss (Figure 3G).

In summary, ChESS-DIA is a facile, robust and scalable assay to measure nuclear protein dynamics across different chromatin environments. Chromatin environments observed in ChESS-DIA correspond to historically-defined functional chromatin fractions defined in the literature and correlate with imaging-based chromatin environments as defined by IF. We applied this method to study the small molecule degrader **dBET1** and provide the first description of the chromatin-wide effects of a PROTAC compound, highlighting a previously unknown interaction between BET bromodomains and histone deacetylase protein complexes. We expect ChESS-DIA to aid provide a new approach to study all nuclear processes that undergo post-transcriptional dynamic regulation.

## Methods

### Cell Culture

K562 and MOLM13 cells were grown in RPMI (GIBCO) supplemented with 10% Fetal Bovine Serum (PAA), sodium pyruvate (GIBCO), L-glutamine (GIBCO), penicillin and streptomycin (GIBCO). **dBET1** (Tocris) was dissolved in DMSO to a concentration of 10mM and cells were treated at a concentration of 1uM for the time indicated.

### Protein Extraction

Nuclear extraction was performed similarly to Dorschner et al [45] by first washing 10^6^ cells with ice cold PBS. All buffers during cell lysis and extraction were supplemented with 1x protease inhibitor solution (Roche). First, cells were resuspended in 0.05% NP-40 dissolved in buffer A (15mM Tris pH 8.0, 15mM NaCl, 60mM KCl, 1mM EDTA pH 8.0, 0.5mM EGTA pH 8). After a 5 minute incubation on ice, nuclei were pelleted at 400xg for 5 minutes. Nuclear extraction was then performed by resuspending the cell pellet first in isotonic Buffer (10mM Tris pH 8.0, 15mM NaCl, 60mM KCl, 1.5 mM EDTA pH 8.0) and incubating at 4°C for 20 minutes. The soluble and insoluble fractions were separated by centrifugation at 500xg for 5 minutes. Next, cells were resuspended in euchromatin extraction buffer (10mM Tris pH 8.0, 250mM NaCl, 1 mM EDTA pH 8.0), incubated at 4°C for 20 minutes and then centrifuged at 1000xg for 5 minutes. Finally, cells were resuspended in heterochromatin extraction buffer (10mM Tris pH 8.0, 600mM NaCl, 1 mM EDTA pH 8.0), incubated at 4°C for 20 minutes and then centrifuged at 20,000xg for 10 minutes. The pellet contains the insoluble protein fraction.

### Protein Preparation and Digestion

Protein samples were supplemented with 0.1% PPS silent surfactant, and the chromatin pellet was sonicated at power 5 for 10s to solubilize proteins. All samples were then boiled at 95°C for 5 minutes, reduced with 5mM DTT at 60°C for 30 minutes and alkylated with 15mM iodoacetic acid (IAA) at 25°C for 30 minutes in the dark. Proteins were then digested with Trypsin (Promega) at 37°C for 2 hours while shaking (1:20 trypsin:protein per sample) and the pH of the sample was subsequently adjusted to <3.0 using HCl. The digested samples were desalted using Oasis MCX 30 mg/60 μm cartridges (Waters) following the manufacturer’s protocol with minor modifications. Briefly, the cartridge was conditioned using 1 mL methanol, 1 mL 10% ammonium hydroxide in water, 2 mL methanol and finally 3 mL 0.1% formic acid in water. The samples were then loaded onto the cartridge and washed with 1 mL 0.1% formic acid in water and 1 mL of 90% acetonitrile in water. The peptides were eluted from the cartridge with 1 mL 10% ammonium hydroxide in methanol which was then removed by evaporation. Peptide samples were resuspended to a concentration of 1ugl in 0.1% formic acid in H2O and stored at -20°C until injected on the mass spectrometer.

### Mass Spectrometry Acquisition

Peptides were separated with a Waters NanoAcquity UPLC and emitted into a Thermo Q-Exactive HF. Pulled tip columns were created from 75 μm inner diameter fused silica capillary in-house using a laser pulling device and packed with 3 μm ReproSil-Pur C18 beads (Dr. Maisch) to 300 mm. Trap columns were created from 150 μm inner diameter fused silica capillary fritted with Kasil on one end and packed with the same C18 beads to 25 mm. Solvent A was 0.1% formic acid in water, while solvent B was 0.1% formic acid in 98% acetonitrile. For each injection, 3 μ! (approximately 1 μg) was loaded and eluted using a 90-minute gradient from 5 to 35% B, followed by a 40-minute washing gradient. For each chromatogram library, the Thermo QExactive HF was configured to acquire six chromatogram library acquisitions with 4 m/z DIA spectra (4 m/z precursor isolation windows at 30,000 resolution, AGC target 1e6, maximum inject time 55 ms) using an overlapping window pattern from narrow mass ranges using window placements optimized by Skyline (i.e. 396.43 to 502.48 m/z, 496.48 to 602.52 m/z, 596.52 to 702.57 m/z, 696.57 to 802.61 m/z, 796.61 to 902.66 m/z, and 896.66 to 1002.70 m/z). See Supplementary Figure 1 or the actual windowing scheme. Two precursor spectra, a wide spectrum (400-1600 m/z at 60,000 resolution) and a narrow spectrum matching the range (i.e. 390-510 m/z, 490-610 m/z, 590-710 m/z, 690-810 m/z, 790-910 m/z, and 890-1010 m/z) using an AGC target of 3e6 and a maximum inject time of 100 ms were interspersed every 18 MS/MS spectra. For quantitative samples, the Thermo Q-Exactive HF was configured to acquire 25x 24 m/z DIA spectra (24 m/z precursor isolation windows at 30,000 resolution, AGC target 1e6, maximum inject time 55 ms) using an overlapping window pattern from 388.43 to 1012.70 m/z using window placements optimized by Skyline. Precursor spectra (385-1015 m/z at 30,000 resolution, AGC target 3e6, maximum inject time 100 ms) were interspersed every 10 MS/MS spectra.

### Mass Spectrometry Data Analysis

Thermo RAW files were converted to .mzML format using the ProteoWizard package (version 3.0.7303) where they were peak picked using vendor libraries and deconvoluted using Prism in “overlap_only” mode. Narrow-window DIA experiments were searched with XCorDIA (manuscript in preparation) using a Human Uniprot protein FASTA file (downloaded 06/10/2017) as both the target and background database. +1H to +4H peptides were detected using default settings (10 ppm precursor and fragment tolerances, trypsin digestion with up to one missed cleavage), assuming fixed cysteine carbamidomethylation. These peptide detections were combined into a chromatogram library, which was used for further processing. EncyclopeDIA was configured to search the wide-window DIA experiments against this chromatogram library with default settings (10 ppm precursor, fragment, and library tolerances, considering both B and Y ions, and trypsin digestion). Both XCorDIA and EncyclopeDIA were configured to use Percolator version 3.1.

### Immunofluoresence

K562 cells were first washed 1x with PBS, resuspended in ice cold buffer-A and permeabilized with 0.025% ice cold NP40 for 4 minutes. The cell solution was then spun down at 500g for 5 minutes, resuspended in ice cold buffer-A and subsequently treated with isotonic, euchromatin and heterochromatin extraction buffers (see “Protein Extraction” methods) for 5 minutes on ice. After the salt treatment, cells were seeded on Poly-L-Lysine coated cover glasses and immediately fixed with 2% PFA for 20 minutes at room temperature. Fixed cells were washed 3x with PBS in preparation for immunofluorescence labeling. Fixed cells were permeabilized with 0.25% PBS-triton for 10 minutes at room temperature, blocked for 1 hour with 3% BSA, and then incubated for 2 hours at room temperature with the relevant primary antibody (1:500 dilution of either anti-rabbit GATA1 (Cell Signaling, D52H6), anti-mouse BRG1 (Santa Cruz, G7), antimouse CRM1 (Santa Cruz, SC74454), and anti-rabbit BRD4 (Bethyl Labs, A301-985A50)), histones (1:500 of either anti-mouse H3K27ac (Active Motif, #39685) or anti-rabbit H3K27ac (Active Motif, #39133)), and 1:500 of anti-goat Lamin B1 (Santa Cruz, SC6216). Subsequently, cells were washed 3×3 minutes with 0.05% PBST, and then incubated for 1 hour with 1:500 dilution of either donkey anti-rabbit Cy3 (#711-166-152, Jackson Labs), donkey anti-mouse AlexaFluor 647 (#715-606-150, Jackson Labs), and donkey anti-goat AlexaFluor594 (A-11058, ThermoFisher). secondary antibodies. Lastly, cells were counterstained with DAPI (100ng/mL) for 10 minutes and washed 3×3 minutes with 0.05% PBST prior to mounting on glass slides using Prolong Gold (Molecular Probes P36930).

2D cell images were acquired using an inverted Nikon Eclipse Ti widefield microscope equipped with a 40x Nikon Plan Apo 0.9 NA air objective and an Andor Zyla 4.2CL10 CMOS camera with a 4.2-megapixel sensor and 6.5μm pixel size (18.8mm diagonal FOV). Acquired images were subject to 3 rounds of iterative blind deconvolution using Autoquant software (version X3.3, Media Cybernetics, NY) to minimize the effect of out-of-focus blurring that is inherent to widefield microscopy optics. Deconvolved images were processed using custom morphometric analysis software to yield numerical estimates for nuclear size and normalized protein content in every cell nucleus.

### Cut & Run

Cut & Run was performed as described previously [44] with minor alterations. 100,000 K562 cells were used per condition. The following antibodies were used:

- BRD2: Active Motif 61797
- BRD3: Active Motif 61489
- BRD4: Bethyl BL-149-2H5
- HDAC1: Active Motif 40967
- HDAC2: Active Motif 39533

After digestion, underwent library preparation with ThruPLEX DNA-seq kit and sequenced on the NextSeq 500 platform (Illumina). Reads were aligned to the human genome (hg38 build) using BWA (version 0.7.12) [48] and peaks called with MACS2 [49] using a q-value cutoff of 0.01.

---

**Supplementary Figure 1:**
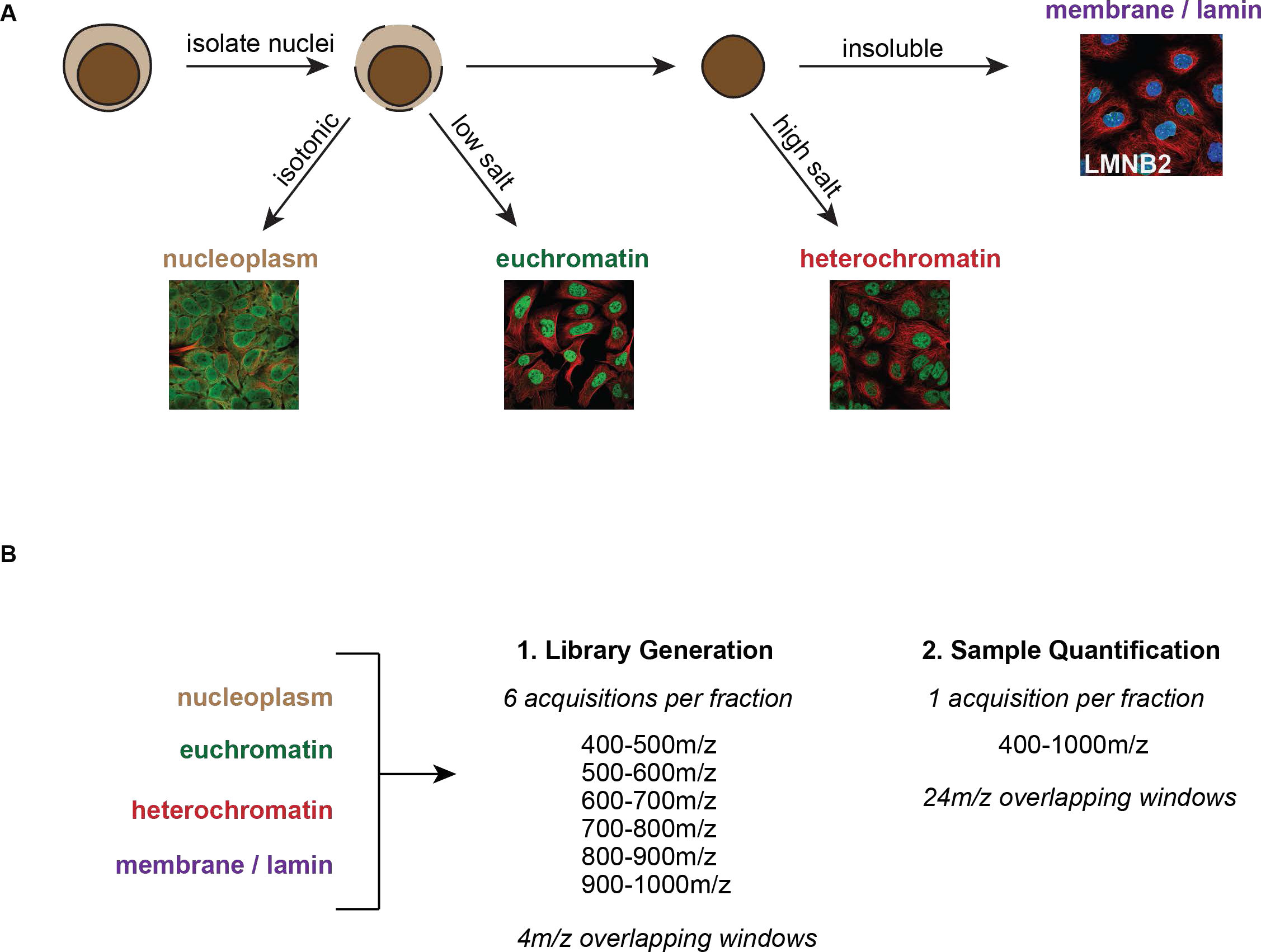
Schematics representing sample preparation and mass spectrometry acquisition for ChESS-DIA. From cells or tissues, nuclei are isolated by treatment with detergent. After isolation, nuclei are then treated with a series of extraction buffers to isolate different chromatin fractions. The fractions are collected and digested for subsequent mass spectrometry. Within an experiment, each fraction is pooled and analyzed by narrow-window DIA to build a chromatogram library. Then, for quantification of individual samples, wide-window DIA is performed on each fraction and analyzed with the EncyclopeDIA software package.

**Supplementary Figure 2:**
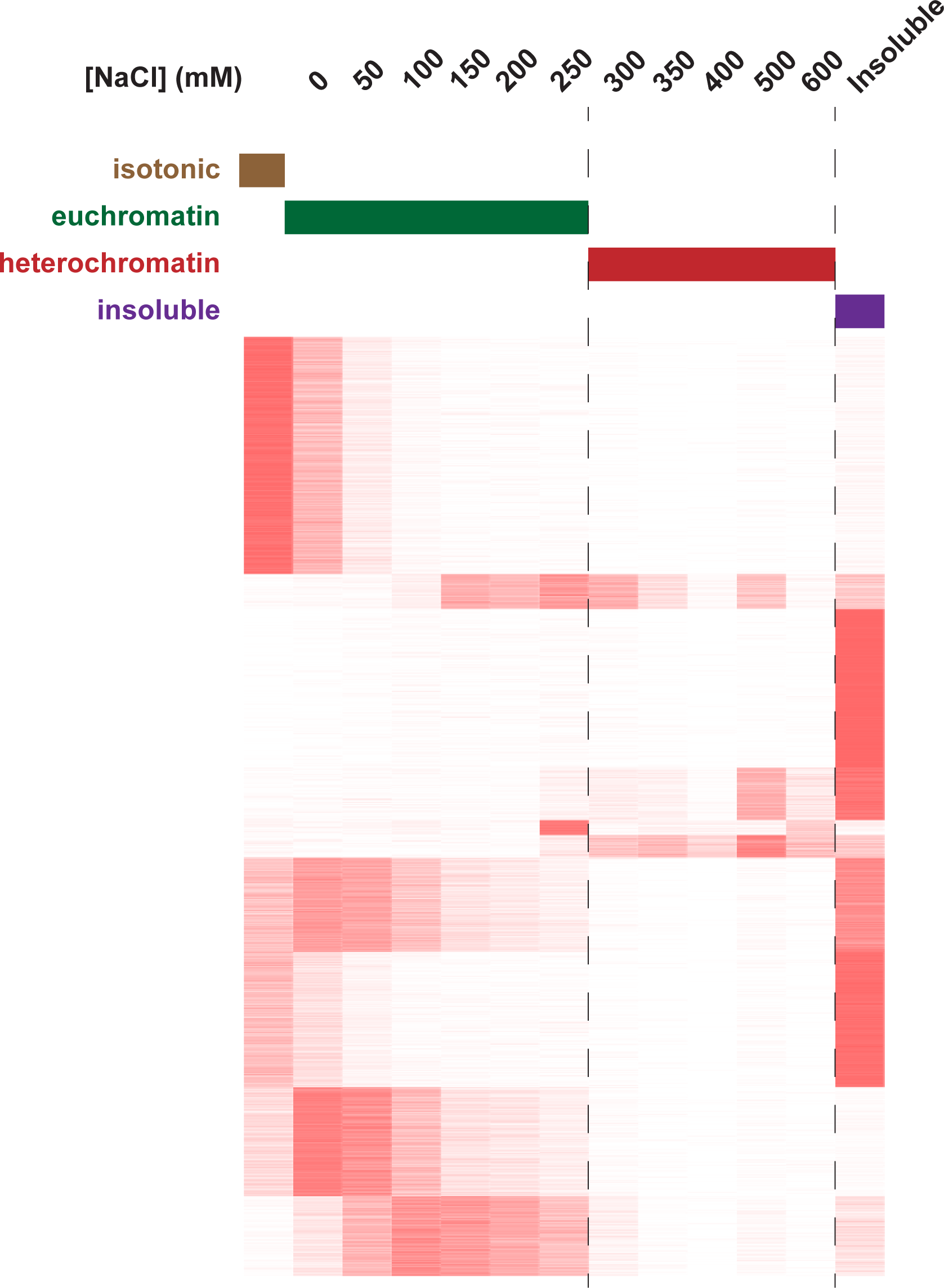
Fine-resolution salt mapping of chromatin. K562 nuclei were isolated and serially extracted with increasing concentrations of salt ranging up to 600mM. Dotted lines indicate the extraction threshold conditions used for all analyses in this study.

**Supplementary Figure 3:**
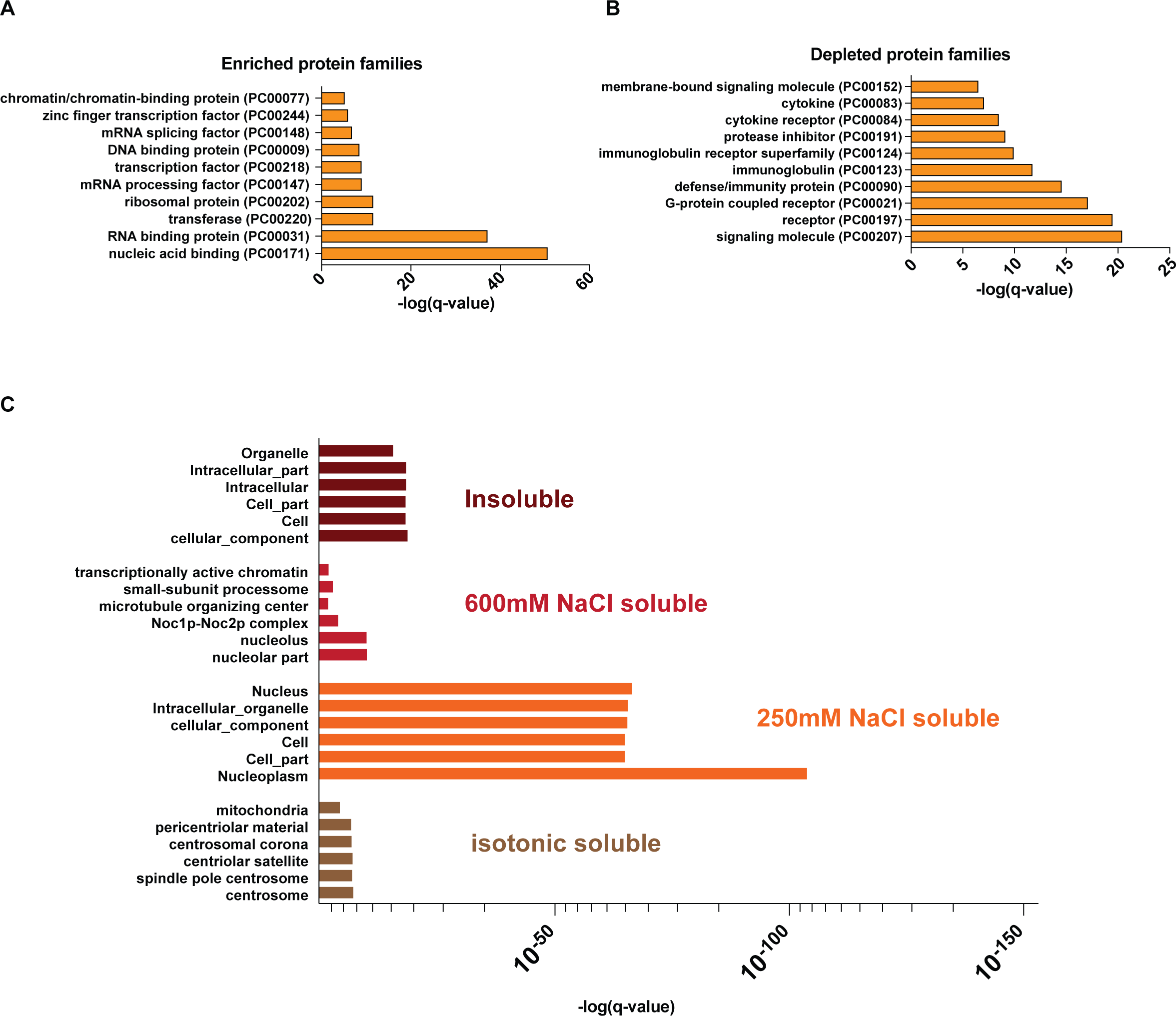
Gene ontology enrichment analysis. (A) PANTHER Gene sets enriched in all proteins isolated by ChESS-DIA in K562 cells(B) PANTHER Gene sets depleted in all proteins isolated by ChESS-DIA in K562 cells (C) Subcellular location gene sets enriched in each fraction measured by ChESS-DIA.

**Supplementary Figure 4:**
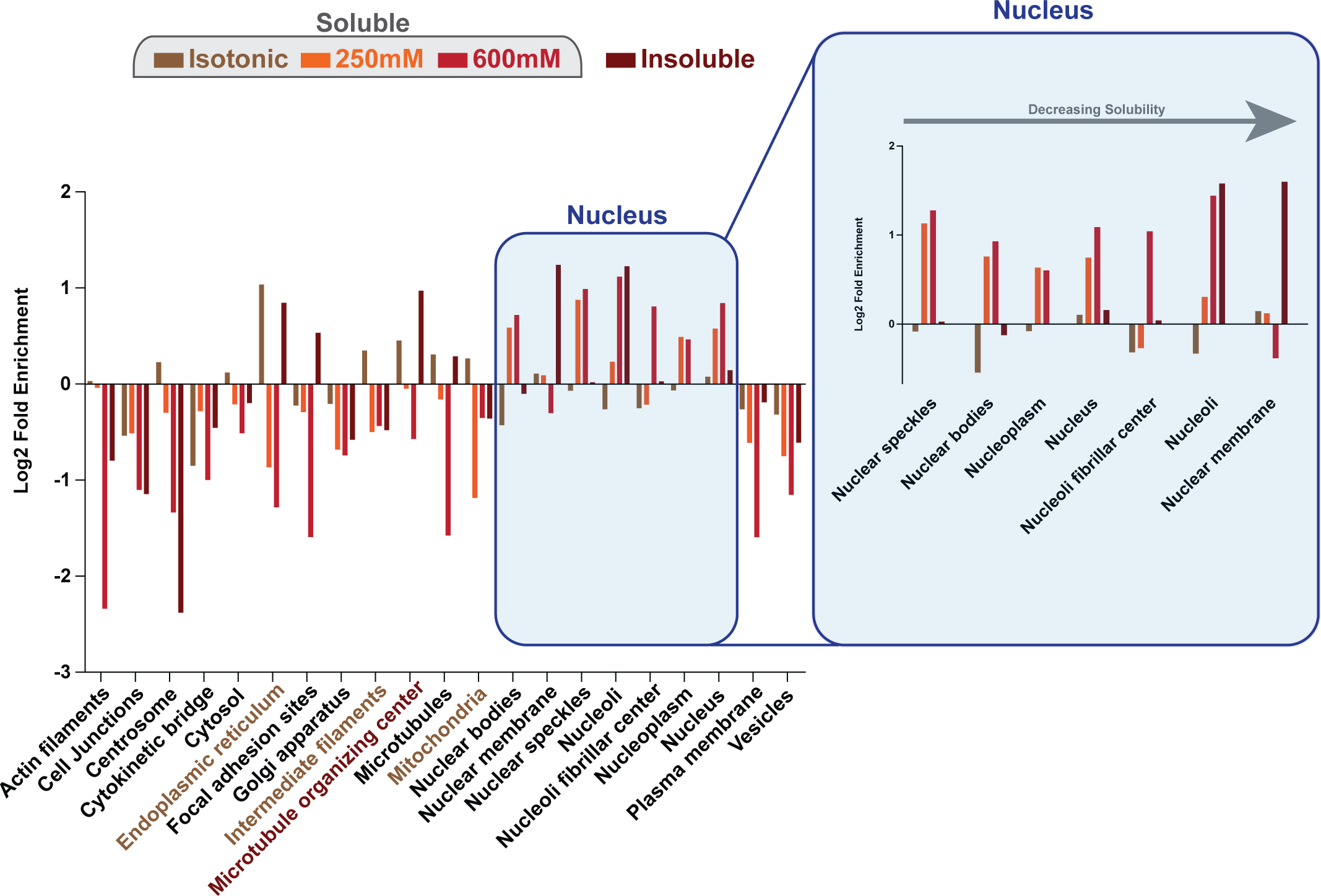
The human protein atlas annotation enrichments across chromatin fractions.

**Supplementary Figure 5:**
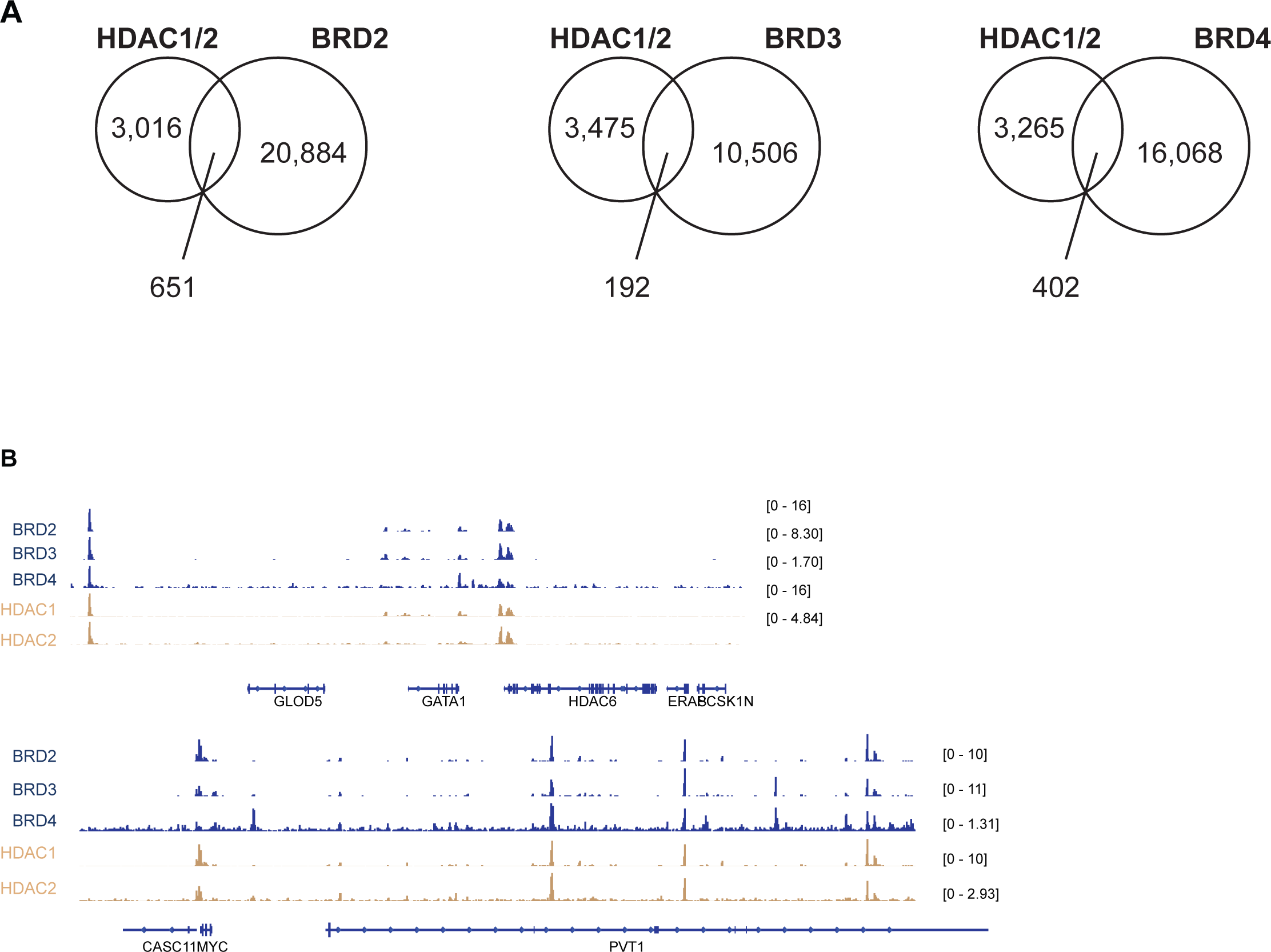
Shared genomic loci between HDAC1/2 and BET bromodomains by Cut&Run. (A) Venn diagrams showing overlapping Cut&Run Peaks between HDAC1/2 shared loci and each BET bromodomain (B) Example gene tracks at MYC and GATA1 gene loci

## References

1. Jackson, D. A., Hassan, A. B., Errington, R. J.& Cook, P. R. Visualization of focal sites of transcription within human nuclei. The EMBO journal 12, 1059–1065 (1993).

2. Lamond, A. I. & Earnshaw, W. C. Structure and function in the nucleus. Science 280, 547–553 (1998).

3. Misteli, T. Protein dynamics: implications for nuclear architecture and gene expression. Science 291, 843–847 (2001).

4. Weintraub, H. & Groudine, M. Chromosomal subunits in active genes have an altered conformation. Science 193, 848–856 (1976).

5. The ENCODE Project Consortium. An Integrated Encyclopedia of DNA Elements in the Human Genome. Nature, 489(7414), 57–74 (2012). http://doi.org/10.1038/nature11247

6. Davis, C. A. et al. The Encyclopedia of DNA elements (ENCODE): data portal update. Nucleic acids research 46, D794–D801 (2017).

7. Wierer, M. & Mann, M. Proteomics to study DNA-bound and chromatin-associated gene regulatory complexes. Hum. Mol. Genet. 25, R106–R114 (2016).

8. Lambert, S. A. et al. The Human Transcription Factors. Cell 172, 650–665 (2018).

9. Novershtern, N. et al. Densely interconnected transcriptional circuits control cell states in human hematopoiesis. Cell 144, 296–309 (2011).

10. Neph, S. et al. Circuitry and Dynamics of Human Transcription Factor Regulatory Networks. Cell 150, 1274–1286 (2012).

11. Shiio Y, Eisenman RN, Yi EC, Donohoe S, Goodlett DR, et al. Quantitative proteomic analysis of chromatin-associated factors. J Am Soc Mass Spectrom 14: 696–703. (2003)

12. Washburn, M. P., Wolters, D. & Yates, J. R. Large-scale analysis of the yeast proteome by multidimensional protein identification technology. Nature Biotechnology 19, 242–247 (2001).

13. Ghaemmaghami, S. et al. Global analysis of protein expression in yeast. Nature 425, 737–741 (2003).

14. Ji, X. et al. Chromatin proteomic profiling reveals novel proteins associated with histone-marked genomic regions. PNAS 201502971–6 (2015).

15. Torrente MP et al. Proteomic Interrogation of Human Chromatin. PLOS One 1–13 (2011).

16. Alajem, A. et al. Differential Association of Chromatin Proteins Identifies BAF60a/SMARCD1 as a Regulator of Embryonic Stem Cell Differentiation. CellReports 10, 2019–2031 (2015).

17. Kustatscher, G. et al. Proteomics of a fuzzy organelle: interphase chromatin. The EMBO Journal 33, 648–664 (2014).

18. Kulej, K. et al. Time-resolved Global and Chromatin Proteomics during Herpes Simplex Virus Type 1 (HSV-1) Infection. Mol Cell Proteomics 16, S92–S107 (2017).

19. Dutta, B., Yan, R., Lim, S. K., Tam, J. P. & Sze, S. K. Quantitative Profiling of Chromatome Dynamics Reveals a Novel Role for HP1BP3 in Hypoxia-induced Oncogenesis. Molecular & Cellular Proteomics: MCP 13, 3236–3249 (2014).

20. Becker, J. S. et al. Genomic and Proteomic Resolution of Heterochromatin and Its Restriction of Alternate Fate Genes. Molecular cell 68, 1023–1037.e15 (2017).

21. Ting, S et al. Peptide-Centric Proteome Analysis: An Alternative Strategy for the Analysis of Tandem Mass Spectrometry Data. 1–23 (2015).

22. Henikoff, S., Henikoff, J. G., Sakai, A., Loeb, G. B. & Ahmad, K. Genome-wide profiling of salt fractions maps physical properties of chromatin. Genome Res. 19, 460–469 (2009).

23. Searle, B. C. et al. Comprehensive peptide quantification for data independent acquisition mass spectrometry using chromatogram libraries. (2018). doi:10.1101/277822

24. Winter, G et al. Phthalimide conjugation as a strategy for in vivo target protein degradation. Science Jun 19;348(6241):1376–81. doi: 10.1126/science.aab1433 (2015).

25. Poss, Z. C., Ebmeier, C. C. & Taatjes, D. J. The Mediator complex and transcription regulation. Critical Reviews in Biochemistry and Molecular Biology 48, 575–608 (2013).

26. Kadoch, C. et al. Proteomic and bioinformatic analysis of mammalian SWI/SNF complexes identifies extensive roles in human malignancy. Nature genetics 45, 592–601 (2013).

27. Dorner, D., Gotzmann, J. & Foisner, R. Nucleoplasmic lamins and their interaction partners, LAP2α, Rb, and BAF, in transcriptional regulation. FEBS Journal 274, 1362–1373 (2007).

28. Morgens, D. W., Deans, R. M., Li, A. & Bassik, M. C. Systematic comparison of CRISPR-Cas9 and RNAi screens for essential genes. Nature biotechnology 34, 634–636 (2016).

29. Ludwig, Anne K., Peng Zhang, and M. C. Cardoso. Modifiers and Readers of DNA Modifications and Their Impact on Genome Structure, Expression, and Stability in Disease. Frontiers in Genetics 7 (2016).

30. Yang, P., Wang, Y. & Macfarlan, T. S. The Role of KRAB-ZFPs in Transposable Element Repression and Mammalian Evolution. Trends in Genetics 33, 871–881 (2017).

31. Thul et al. A subcellular map of the human proteome. Science (2017).

32. Barrales, Ramón Ramos et al. Control of Heterochromatin Localization and Silencing by the Nuclear Membrane Protein Lem2. Genes & Development 30.2 (2016).

33. Neklesa, T. K., Winkler, J. D. & Crews, C. M. Targeted protein degradation by PROTACs. Pharmacology & Therapeutics 174, 138–144 (2017).

34. Filippakopoulos, P. et al. Histone Recognition and Large-Scale Structural Analysis of the Human Bromodomain Family. Cell 149, 214–231 (2012).

35. Roe, Jae-Seok et al. BET Bromodomain Inhibition Suppresses the Functional Output of Hematopoietic Transcription Factors in Acute Myeloid Leukemia. Molecular cell 58.6 (2015).

36. Lovén, J. et al. Selective Inhibition of Tumor Oncogenes by Disruption of SuperEnhancers. Cell 153, 320–334 (2013).

37. Aftimos, P. G. et al. Phase I first-in-man trial of a novel bromodomain and extra-terminal domain (BET) inhibitor (BI 894999) in patients (Pts) with advanced solid tumors. J. Clin. Oncol. 35, 2504–2504 (2017).

38. Hsu, Sarah C. et al. The BET Protein BRD2 Cooperates with CTCF to Enforce Transcriptional and Architectural Boundaries. Molecular cell 66.1 (2017).

39. Decker, T.-M. et al. Transcriptome analysis of dominant-negative Brd4 mutants identifies Brd4-specific target genes of small molecule inhibitor JQ1. Scientific Reports 7, 1684 (2017).

40. Roberts, T. C. et al. BRD3 and BRD4 BET Bromodomain Proteins Differentially Regulate Skeletal Myogenesis. Scientific Reports 7, 6153 (2017).

41. Brown, JD et al. NF-kB directs dynamic super enhancer formation in inflammation and atherogenesis. Mol Cell 23; 56(2): 219–231 (2014).

42. Bradner JE et al. Transcriptional Addiction in Cancer. Cell 629–643 (2017)

43. Basta, J. & Rauchman, M. The nucleosome remodeling and deacetylase complex in development and disease. Translational Research 165, 36–47 (2015).

44. Skene PJ, Henikoff S. An efficient targeted nuclease strategy for high-resolution mapping of DNA binding sites. Elife, 21856. (2018).

45. Dorschner, M. O. et al. High-throughput localization of functional elements by quantitative chromatin profiling. Nature Methods 1, 219 (2004).

46. Ting, Y. S. et al. PECAN: Library Free Peptide Detection for Data-Independent Acquisition Tandem Mass Spectrometry Data. Nature Methods 14, 903–908 (2017).

47. Käll, L., Canterbury, J. D., Weston, J., Noble, W. S. & MacCoss, M. J. Semi-supervised learning for peptide identification from shotgun proteomics datasets. Nature Methods 4, 923 (2007).

48. Li, H. & Durbin, R. Fast and accurate short read alignment with Burrows-Wheeler transform. Bioinformatics 25, 1754–1760 (2009).

49. Zhang, Y. et al. Model-based Analysis of ChIP-Seq (MACS). Genome Biology 9, R137–R137 (2008).

